# The Brain Imaging Data Structure, a new format for organizing and describing outputs of neuroimaging experiments

**DOI:** 10.1101/034561

**Authors:** Krzysztof J. Gorgolewski, Tibor Auer, Vince D. Calhoun, R. Cameron Craddock, Samir Das, Eugene P. Duff, Guillaume Flandin, Satrajit S. Ghosh, Tristan Glatard, Yaroslav O. Halchenko, Daniel A. Handwerker, Michael Hanke, David Keator, Xiangrui Li, Zachary Michael, Camille Maumet, B. Nolan Nichols, Thomas E. Nichols, John Pellman, Jean-Baptiste Poline, Ariel Rokem, Gunnar Schaefer, Vanessa Sochat, William Triplett, Jessica A. Turner, Gaël Varoquaux, Russell A. Poldrack

## Abstract

The development of magnetic resonance imaging (MRI) techniques has defined modern neuroimaging. Since its inception, tens of thousands of studies using techniques such as functional MRI and diffusion weighted imaging have allowed for the non-invasive study of the brain. Despite the fact that MRI is routinely used to obtain data for neuroscience research, there has been no widely adopted standard for organizing and describing the data collected in an imaging experiment. This renders sharing and reusing data (within or between labs) difficult if not impossible and unnecessarily complicates the application of automatic pipelines and quality assurance protocols. To solve this problem, we have developed the Brain Imaging Data Structure (BIDS), a standard for organizing and describing MRI datasets. The BIDS standard uses file formats compatible with existing software, unifies the majority of practices already common in the field, and captures the metadata necessary for most common data processing operations.

---

Neuroimaging, the study of the brain with medical-imaging devices such as magnetic resonance scanners, is our number one source of quantitative data on brain structure and function. Based on the volume of publication^1^, tens of thousands subjects are scanned for research purposes each year. Each study results in complex data involving many files in different formats ranging from simple text files to multidimensional image data, which can be arranged in many different ways. Indeed, a study is typically comprised of multiple imaging protocols and often multiple groups of subjects. To date there has been no consensus about how to organize and share these data, leading researchers, even those working within the same lab, to arrange their data in different and idiosyncratic ways. Lack of consensus leads to misunderstanding and time wasted on rearranging data or rewriting scripts that expect particular file formats and organization, as well as a possible cause for errors. Adoption of a common standard to describe data and its organization on disk can provide multiple benefits:

- **Minimized curation:** Common standards make it possible for researchers who were not directly involved in data collection to understand and work with the data. This is particularly important to ensure that data remain accessible and usable by different researchers over time in the following instances:

a) within a laboratory over time,
b) between labs facilitating collaboration and making combining data in multi-center studies easier and less ambiguous,
c) between public databases (i.e. OpenfMRI) allowing for the quick ingestion of big data organized according to a common scheme.
- **Error reduction:** Errors attributed to the misunderstanding of the meaning of a given datum (e.g., when variable names are not explicitly stated in the data file and standardized across files).
- **Optimized usage of data analysis software** is made possible when the metadata necessary for analysis (i.e. details of the task or imaging protocol) are easily accessible in a standardized and machine-readable way. This enables the application of completely automated analysis workflows, which greatly enhances reproducibility and efficiency.
- **Development of automated tools** for verifying the consistency and completeness of datasets is realized. Such tools make it easier to spot missing metadata that limit how the data could be analyzed in the future.

Previous approaches to neuroimaging data management typically involved complex data management systems^2–8^. However, the challenges associated with installing and maintaining an additional software application, and interacting with one’s data primarily through that application, may outweigh the benefits for smaller labs with limited technical resources^9^. In addition, the vast majority of data analysis software requires access to files stored on a hard drive, which is only directly supported by some neuroimaging data management systems. This leads to the need for exporting datasets to a filesystem as the first step of any analysis^10^.

The goal of previous databasing approaches has been to efficiently store and manage data rather than creating a format for describing and standardizing it. In contrast, the XML-based Clinical Experiment Data Exchange schema (XCEDE)^11^ attempted to provide a standard for describing results of clinical, including neuroimaging, experiments (independent of any particular databasing system). The approach used by XCEDE employs the extensible Markup Language (XML)^12^ to provide a hierarchical description of a dataset. This description includes location of every data file along with metadata. Due to the fact that location of files is decoupled from their purpose, XCEDE supports any arbitrary arrangement of files on the hard drive (or even remote locations). In addition, it does not provide any recommendation on the choice of the file format that the imaging data should be stored in (which puts burden of data conversion on the shoulders of tool developers). Unfortunately, the XCEDE format was not widely

adopted, although a number of useful tools were developed^13–15^. We suspect that the combination of extensive use of XML (which is hard to use for scientists without informatics expertise), lack of specification of the file format details, as well as relatively limited support in data analysis packages all may have contributed to the low adoption rate.

In a similar fashion to XCEDE, the OpenfMRI database^16^ introduced a dataset description format to fulfill the needs of data curation and dissemination. It relies heavily on specific file naming schemes and paths to convey the functions of files, which allows the application of automated analysis workflows for the entire processing stream. The use of specific filenames and paths can be initially viewed as a limitation in contrast to XCEDE, but it makes it much easier to write software to analyze the data since it does not require consulting additional files (XML descriptions) to understand the purpose of a particular file. In addition, the OpenfMRI standard uses the Gzip compressed version of the Neuroinformatics Informatics Technology Initiative (NIfTI) format^17^ to achieve a balance of data analysis software compatibility and file size. This avoids the need to convert between file formats as may have been necessary with XCEDE since it did not specify a particular file format (given that many software tools are limited in the formats that they can read). An important limitation of the OpenfMRI standard is that it had no explicit support for a number of important data types including physiological recordings, diffusion weighted imaging, or field maps, and also had no formal scheme to accommodate longitudinal studies with multiple visits. Despite these limitations and the fact that the OpenfMRI standard was designed to fulfill the needs of one particular repository, it has provided a unified and simple way to organize and describe data. This led to it being adopted (with some modifications) as an internal standard for organizing data in a number of laboratories as well as support by Nipype workflow engine^18^.

Dataset description can also be considered part of provenance. W3C PROV standard defines provenance as “information about entities, activities, and people involved in producing a piece of data or thing, which can be used to form assessments about its quality, reliability or trustworthiness”(https://www.w3.org/TR/prov-overview/). Considering this definition metadata describing for example scanner field strength used to obtain MRI data fall into this category. However, most neuroimaging literature discussing provenance focuses on recording entities and activities interacting with data after it was acquired (for example version of software used to perform spatial smoothing)^19,20^. An exception to this trend is work of Mackenzie-Graham et al. proposing a provenance scheme that included information about data acquisition parameters^21^. However, because this scheme was mainly focused on data processing provenance it only included a few acquisition parameters (such as repetition time, pulse sequence, coil type etc.) and did not include paradigm details and other metadata crucial for analyzing task fMRI or diffusion data.

When developing the Brain Imaging Data Structure (BIDS) standard, proven parts from the aforementioned standards were combined with common laboratory practices to maximize ease of use and adoption. Common practices included encoding the purpose of a file in its filename and reusing already existing and widely recognized file formats (NIfTI, JavaScript Object Notation [JSON], and Tab Separated Value [TSV] text files). The process of defining this standard involved consultations with leading scientists in the field, public calls for comments, and most importantly the generation of example BIDS compatible versions of publicly available MRI datasets. The resulting specification is intentionally based on simple file formats (often text-based) and folder structures. This is done to reflect common lab practices in the community and to make it accessible to a wide range of scientists with limited technical backgrounds. Additional metadata (e.g. acquisition details) are stored in JSON files^22^. JSON is arguably easier to write and comprehend than XML^23^, is widely supported by major programming languages, and can be linked to formal ontologies (e.g., Cognitive Atlas^24^, Cognitive Paradigm Ontology^25^, and NIDM^26^) via JSON-LD^27^.

## Results

The following standard describes a way of arranging data (see Figure 1) and specifying metadata for a subset of neuroimaging experiments. It follows a simple but carefully defined terminology. The filenames are formed with a series of key-values and end with a file type, where keys and file types are predefined and values are chosen by the user. Some aspects of the standard are mandatory. For example, each dataset needs to have at least one subject directory. Some aspects are regulated but optional. For example, T1-weighted scans do not need to be included, but when they are available they should be saved under a particular file name pattern specified in the standard. The standard provides data dictionaries and strict naming conventions for structural (T1w, T2w etc.), diffusion, and functional MRI data as well as accompanying behavioural and physiological data. In addition clear definitions of terms used in TSV and JSON files are provided together with links to DICOM, Cognitive Atlas^24^. and Cognitive Paradigm^25^ ontologies.

**Figure 1.**
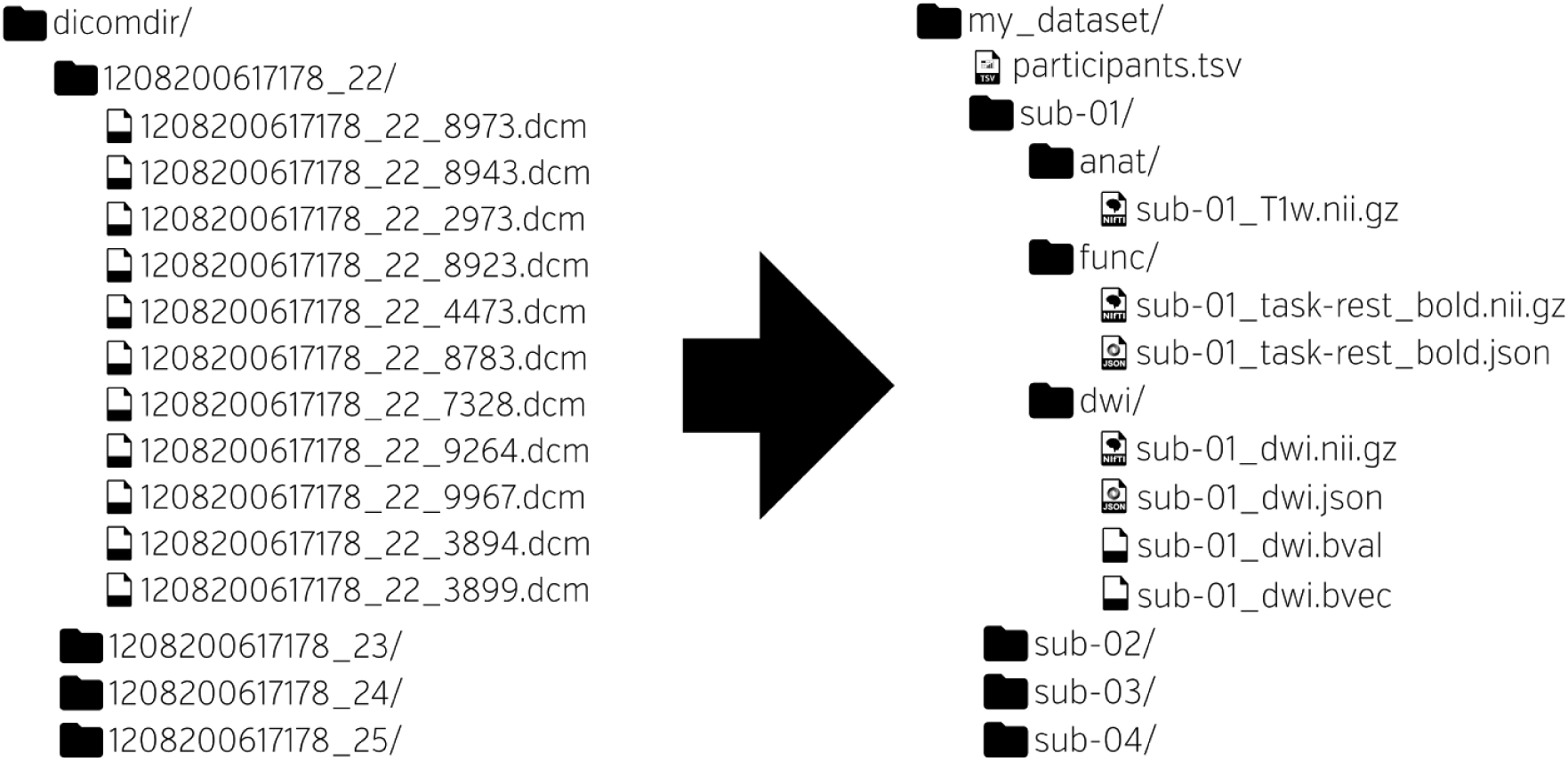
BIDS is a protocol for standardizing and describing outputs of neuroimaging experiments (left) in a way that is intuitive to understand and easy to use with existing analysis tools (right).

This standard aspires to describe a majority of datasets, but acknowledges that there will be cases that do not fit the present version (1.0.0) of BIDS. In such cases one can include additional files and subfolders to the existing folder structure following a set of general naming guidelines and common sense. For example, one may want to include eye tracking data (BIDS does not cover this type of data yet). A sensible place to put it is next to the continuous recording file with the same naming scheme but different extensions. To make sure such additions are not accidental the provided validator raises a warning for all files that do not fit the specification. The solutions will vary from case to case and publicly available datasets will be periodically reviewed to include common data types in the future releases of the BIDS specification.

### Raw vs. derived data

BIDS in its current form is designed to standardize (convert to a common file format) and describe raw data. During analysis, such data will be processed and intermediate as well as final results will be saved. Derivatives of the raw data, should be kept separate from the raw data. This clearly separates raw from processed data, makes sharing of raw data easier, and prevents accidental changes to the raw data. Even though BIDS specification currently does not contain a particular naming scheme for different data derivatives (correlation maps, brain masks, contrasts maps, etc.) we recommend keeping them in a separate “derivatives” folder with a similar folder structure as presented below for the raw data. For example: derivatives/sub-01/ses-pre/mask.nii.gz. In the future releases of BIDS we plan to provide more detailed recommendations on how to organize and describe various data derivatives.

### The Inheritance Principle

Any metadata file (e.g., files ending with: .json, .bvec, _events.tsv, _physio.tsv.gz, and _stim.tsv) may be defined at one of four levels (in hierarchical order): MRI acquisition, session, subject, or dataset. Values from the top level are inherited by all lower levels unless they are overridden by a file at the lower level. For example, /task-nback_bold.json may be specified at the dataset level to set Time of Repetition (TR) for all subjects, sessions and runs. If one of the runs has a different TR than the one specified in the dataset level file, a /sub-<subject_id>/sub-<subject_id>_task-nback_bold.json file can be used to specify the TR for that specific run.

### File Formats

#### Imaging files

Since BIDS is aimed at facilitating data sharing as well as analysis the file format for storing imaging data was selected based on support from various neuroimaging data analysis packages. We have chosen the NIfTI file format because it is the largest common denominator across neuroimaging software. However, since it offers limited support for the various image acquisition parameters available in DICOM or other scanner specific files, the BIDS standard requires users to provide additional meta information in a sidecar JSON file (with the same filename as the .nii.gz file, but with a .json extension - see section”Key/value files” for more information). BIDS standard specifies a carefully selected set of fields together with their definitions which extends the standard DICOM ontology with terms that are crucial for data analysis such as the polarity of phase encoding direction or slice timing (which traditionally have been recorded in inconsistent ways across scanner manufacturers and are not part of the DICOM ontology). In addition to terms specified in BIDS we encourage users to include other information extracted from DICOMs (including private manufacturer fields) during the conversion process so no metadata would be lost. Extraction of a minimal set of BIDS compatible metadata can be performed using dcm2niix (https://www.nitrc.org/proiects/dcm2nii/) and dicm2nii (http://www.mathworks.com/matlabcentral/fileexchange/42997) DICOM to NIfTI converters. A provided validator (https://github.com/INCF/bids-validator) will check completeness of provided metadata and look for conflicts between the JSON file and the data recorded in the NIfTI header.

#### Tabular files

Meta-data most naturally stored as an array are stored in tab-delimited value (TSV) files, similar to comma-separated value (CSV) files where commas are replaced by tabs. A header line is generally required naming each column and, depending on the use, some specific variable names are required (see the full specification for details). String values containing tabs should be escaped using double quotes.

Missing values should be coded as “n/a”.

#### Key/value files (dictionaries)

JSON files will be used for storing key/value pairs, with the key names following a fixed dictionary in the specification. Extensive documentation of the JSON format can be found at http://json.org. Several editors have built-in support for JSON syntax highlighting that aids manual creation and editing of such files. An online editor for JSON with built-in validation is available at http://jsoneditoronline.org. JSON files need to be encoded in ASCII or UTF-8. The order of keys is arbitrary and should does not convey any meaning.

### Required, recommended, and optional metadata

To maximize adoption and flexibility of the BIDS standard only a small subset of metadata fields and files is required (compulsory). The decision which metadata fields and files are required was based on the minimal metadata needed to perform standard basic analyses on each type of data. For anatomical scans only specifying the type (T1 weighted, T2 weighted, T1 map etc. see Section 8.3 in Supp. File 1) is required. For functional scans (fMRI), the researcher is required to specify task name (which could be “rest” in for so-called resting-state scans), repetition time (in seconds) and timing and duration of all events (stimuli and/or responses, unless the subject was not performing any task; for more details see Section 8.4 in Supp. File 1). For diffusion weighted imaging the required metadata is limited to b-values (in the form of .bval files) and diffusion gradient tables (in the form of .bvec files; for more information see Section 8.8 in Supp. File 1). Different types of fieldmaps also include a set of corresponding required fields (see Section 8.9 in Supp. File 1). Similarly when including physiological (breathing or cardiac) or other continuous recordings the researcher is required to specify a start time (relative to the beginning of image acquisition) and sampling frequency (for more details see Section 8.6 in Supp. File 1). When a required file or metadata field is missing the BIDS Validator will report an error.

In addition to those mandatory pieces of metadata, the BIDS standard strongly recommends inclusion of other metadata that are crucial for performing some additional types of analyses. Those include, but are not limited to slice timing (necessary for slice timing correction), phase encoding direction, effective echo spacing, and echo time (required for performing field unwarping). When a recommended piece of metadata is missing, the BIDS Validator will report a warning.

Finally, the BIDS specification also defines a large set of metadata fields that are optional. Those include information that is not crucial for any particular data analysis method, but can be useful when trying to understand the nature of the data or combining data from multiple sources. Those fields include, but are not limited to scanner manufacturer, scanner software version, head coil name, instructions given before the task, multiband acceleration factor etc. In addition the researcher can extend the metadata dictionaries with their own keys (as long as they do not collide with those already defined in BIDS specification) to include additional information.

### Creating a BIDS compatible dataset

The process of creating a BIDS compatible dataset can be split into several steps. In the following section we will present this procedure using a dataset acquired at UCLA by Jessica Cohen as a part of her Ph.D. research ^28^. This dataset includes anatomical, diffusion and task fMRI data and is available (in BIDS format) in OpenfMRI repository under the accession number ds000009.

#### Step 1: Convert DICOM files to NIfTI

This dataset has been acquired using an MRI scanner that outputs DICOM files (Siemens Trio) so we can use a DICOM to NIfTI converter such as dcm2niix. This particular converter supports BIDS -- it normalizes idiosyncrasies of different scanner manufacturers that are not standardized by DICOM, and outputs a BIDS compatible JSON with most of the required and recommended metadata (such as repetition time, slice timing, and phase encoding direction).

#### Step 2: Create folder structure, rename and copy NIfTI files

BIDS relies heavily on a particular folder structure and naming scheme of files. We begin creating the folder structure, by creating one subfolder for each of the 29 subjects named “sub-01”, “sub-02”, “sub-03” etc. Inside each of the subject subfolders we create three subfolders: “anat” (for anatomical scans), “dwi” (for diffusion scans), and “func” (for task fMRI). Those names are not arbitrary and must follow the BIDS specification (see Supp. File 1). This dataset includes two anatomical scans per subject: high-resolution T1 weighted and in-plane T2 weighted. They need to be renamed to “sub-01_T1w.nii.gz“ and “sub-01_inplaneT2.nii.gz” (respectively) and moved to the “anat” subfolder. This operation has to be repeated for all subjects. Along the .nii.gz files .json files (with the same body of the file name) should be also moved.

Similarly we move the diffusion files into the “dwi” folder. The naming scheme is analogous “sub-01_dwi.nii.gz”. In addition to .json files we also move the .bvec and .bval files containing gradient information produced by dcm2niix. Finally we follow suit with the task fMRI files. This dataset includes four different tasks with the following names: stop-signal, Balloon analog risk task (BART), discounting, and emotion regulation, which we label as “stopsignal”, “bart”, “discounting” and “emotionregulation” correspondingly. The naming scheme for functional is “sub-01_task-stopsignal_bold.nii.gz” (where “01” is replaced by corresponding subject label for the other subjects and “stopsignal” is replaced by corresponding task label for the other tasks).

#### Step 3: Add remaining data

In addition to imaging data and metadata we also need to provide details of the experimental paradigm for the task fMRI data. This is done by creating a tab-delimited text file following the naming scheme of “sub-01_task-stopsignal_events.tsv” for each of the .nii.gz files. These files includes two compulsory columns: “onset” and “duration” (both in seconds) and any number of other arbitrary named columns to categorize and describe events (both stimuli and responses) recorded during the experiments. In case of this task we will add columns describing reaction time (in seconds), trial type (go or stop), subject response, response correctness, and trial outcome.

On top of the experimental paradigm information we also have some demographic information about the participants of the study such as age and sex. This data should be saved in a text file called “participants.tsv” in the root of the dataset directory. This file has one compulsory column: “participant_id” (for example “sub-01”, “sub-02”) and can include any number of other arbitrarily named columns describing participants. Optionally a “participants.json” file can be provided with description of each column and links to external ontologies (see Section 4.2 in Supp. File 1).

#### Step 4: Add missing metadata

All of the metadata in .json files were so far obtained using the dcm2niix converter. In addition to this we need to provide the name of each fMRI task. Optionally we can add information about task instructions and description as well as link the tasks an external ontology such as Cognitive Atlas or Cognitive Paradigm Ontology. Metadata organization can also be simplified using the inheritance rule: Metadata fields common across all subjects can be specified in one JSON file in the root of the directory instead of being repeated for each subject (see Section 3.5 in Supp. File 1 for details). Finally we need to create a dataset_description.json file with fields that include the name and description of the dataset as well as the version of BIDS standard used. This file can also be used to list authors and ways to reference the dataset (see Section 8.1 in Supp. File 1).

#### Step 5: Validate the dataset

Once the dataset is assembled, the BIDS Validator can be used to check if any of the required or recommended metadata are missing. In addition the validator has built in heuristics to spot incorrect definitions of missing values (for example “NA” instead of “n/a”), use of wrong units (milliseconds instead of seconds), missing scans and inconsistent scanning parameters across subjects. The validator works in the Chrome web browser with no need to install additional software, and performs the validation on the client side (i.e., no data are uploaded or shared) so it is suitable for sensitive datasets that are not intended for public sharing.

Any BIDS compatible dataset can be readily fed into MRIQC or QAP toolboxes (see Adoption) that calculate quality measures. Thanks to formal structure of BIDS no additional metadata are required as an input. Outputs of those quality analyses can be included along with the dataset (see Section 3.4 in Supp. File 1). Similarly BIDS2ISATab tool can be used (again with no need to specify additional metadata) to generate metadata summary files that are required for publishing with Scientific Data and GigaScience journals.

### Adoption

Despite its relatively young age BIDS has been already adopted by the OpenfMRI repository^16^. Since the switch to the new standard in December 2015, thirteen new BIDS compatible datasets have been published. In addition several software packages added support for BIDS: SciTran (database)^29^, Quality Assurance Protocol (QA toolbox - https://github.com/preprocessed-connectomes-project/quality-assessment-protocol), MRIQC (QA toolbox - https://github.com/poldracklab/mriqc), and automatic analysis (workflow toolbox)^30^ have added BIDS support. In addition, a number of tools have been developed to help working with BIDS datasets. Those include: bids-validator (a validation tool -https://github.com/INCF/bids-validator), openfmri2bids (OpenfMRI convention to BIDS converter - https://github.com/INCF/openfmri2bids), BIDSto3col (FSL modelling helper tool - https://github.com/INCF/bidsutils/tree/master/BIDSto3col), and BIDS2ISATab (tool for extracting ISA-Tab^31^ compatible metadata from BIDS datasets to improve speed and accuracy of data curation journals using this standard - https://github.com/INCF/BIDS2ISATab).

## Discussion

Since BIDS was designed to maximize adoption, it heavily relies on established file formats such as NIfTI and bvec/bval (see the protocol for details). This decision was made because those file formats are widely supported by neuroimaging software. Using other file formats (such as DICOM which is closer to the scanner output or HDF5 which is much more flexible and allows for storing all metadata) would result in a more concise and robust data structure, albeit at the cost of additional software development necessary to adapt existing software to the new file format. Storing metadata in JSON files has advantages of accessibility, but can be error prone because data and metadata do not live in the same file. In future revisions of BIDS we will explore the possibility of storing metadata as a JSON text extension of the NIfTI header.

While we chose NIfTI to store neuroimaging data due to its popularity, we also recognize that specific tools or communities use other neuroimaging file formats such as MINC or NRRD for both technical and historical reasons. Different flavors of BIDS can be designed to support such formats, which would additionally require (1) identification of the metadata fields that should be included in the sidecar JSON file (as opposed to the data file headers), and (2) modifying the validator to read the new file format and check for required and optional metadata. For example, the MINC community (represented in this work by authors SD and TG) is currently working on an mBIDS specification that is based on BIDS, but uses MINC file format instead of NIfTI. Even though having multiple flavors of one standard can be problematic and confusing for software developers, those flavors are also necessary to meet the specific needs of some communities. To avoid confusion, any future derivatives of the BIDS specification that are not compatible with the original should be clearly marked in the dataset metadata. At the same time we acknowledge that NIfTI file format is far from perfect and it quite likely that it will be replaced by a solution more capable of random access to large compressed blocks of data and with built-in extensible metadata storage. Through initiatives such as mBIDS we can decouple file organization from a particular file format used for storing imaging data.

The current release of BIDS does not include support for Electroencephalography (EEG) and Magnetoencephalography (MEG) data, because, at present, there is no single commonly accepted data exchange file format for such data (akin to NIfTI in neuroimaging). However, we plan to extend the standard with support for EEG/MEG in a future release. Similarly the current version of the standard does not cover Positron Emission Tomography (PET), Arterial Spin Labeling (ASL) and spectroscopy, but those extensions will be considered in the future.

Our major focus in the near future will be on extending the software ecosystem supporting BIDS to provide incentives for researchers to use it. Work is underway to include BIDS support within heudiconv (data organizer), PyMVPA (statistical learning toolbox), automatic analysis (framework for analysing multimodal datasets), C-PAC (resting state analysis toolbox), and CBRAIN (data analysis platform). Furthermore, XNAT, COINS and LORIS databases are planning to support BIDS as an export and/or import option. We also plan to build tools to facilitate conversion to the NIMH Data Archive.

It is also important to acknowledge that formal, machine readable descriptions of datasets such as BIDS can only complement, but not replace free form descriptions written in prose such as data papers or data descriptors ^32^.

Such unconstrained format is not only capable to capture motivation behind acquiring a particular dataset or describe in detail experimental design of a particular behavioural paradigm, but also can be easily turned into an academic paper providing an important incentive for data sharing. Adoption of BIDS in data papers describing neuroimaging data can increase their value, because BIDS will make it easier to assess completeness and internal validity of a dataset and make it easier to reuse.

This article serves only as an introduction to the BIDS standard - the complete version of the the specification is available as supplementary material. In addition, for example datasets, a list of resources, and pointers on how to give feedback on future releases please visit http://bids.neuroimaging.io.

## Methods

Work on the Brain Imaging Data Structure began at a meeting of the INCF Neuroimaging Data Sharing Task Force (wiki.incf.org/mediawiki/index.php/Neuroimaging_Task_Force) held at Stanford University on January 27-30th 2015. While a flexible solution using the PROV W3C model (http://www.w3.org/TR/prov-overview/) was first investigated, it was acknowledged that this technology would be only viable if tools were in place to write the associated metadata. Since experimental data are obtained from multiple tools, a solution accessible to most neuroimaging researchers was designed. An initial draft was heavily inspired by the data structure used by the OpenfMRI database, but soon evolved beyond backward compatibility. After the initial draft was formed, a series of discussions and public calls for feedback were conducted. Feedback was solicited over Twitter, by presenting BIDS and distributing informational pamphlets at conferences (INCF, SfN), as well as by sending emails to SPM, FSL, Freesurfer, MRTrix, Slicer, Nipy, and HCP mailing lists (reaching over 5,000 researchers). Further refinement of the standard was facilitated by a meeting held during the OHBM conference in Honolulu in June 2015. The discussion over the standard involved domain researchers, computer scientists, MRI physicists, methods developers (FSL, SPM, Slicer, Nipy, PyMVPA, C-PAC, nilearn and aa), data curators (OpenfMRI, FCP/INDI, HCP, NKI, SchizConnect, ABIDE, DataLad, and BIRN) and database developers (COINS, LORIS, XNAT, NiDB and SciTran). The first Release Candidate was published on September 21st 2015 along with 22 example datasets, online and command line validation tools (https://github.com/INCF/bids-validator), and a converter from OpenfMRI standard (https://github.com/INCF/openfmri2bids). The standard became official (version 1.0.0) with the publication of this manuscript and we expect to update and extend it through future releases (see Supp. File 1). We encourage everyone to provide feedback on the standard as well as suggestion for new features and support for more data types. Proposed changes will be discussed publicly trying to accommodate the needs of the community. To facilitate this process we have created the http://bids.neuroimaging.io website.

## ACKNOWLEDGEMENTS

This work has been supported by the International Neuroinformatics Coordinating Facility (INCF) and by the Laura and John Arnold Foundation. VDC has been supported by in part by NIH P20GM103472. TEN and CM have been supported by the Wellcome Trust. NN has been supported by NIH NIAAA [1 U01 AA021697]. JBP, BNN, and RAP have been supported by NIH NIAAA OD [1 U01 AA021697-04S1]. DAH was supported by the Intramural Research Program of the NIMH. MH was supported by the German federal state of Sachsen-Anhalt and the European Regional Development Fund (ERDF), project: Center for Behavioral Brain Sciences (CBBS). T.A. was supported by the Medical Research Council (United Kingdom) [MC-A060-53144]. YOH has been supported by in part by NSF [1429999].

